# Molecular characterization of unusual G10P[33], G6P[14] genomic constellations and evidence of zooanthroponosis in bovines

**DOI:** 10.1101/2020.02.25.965731

**Authors:** P.M Sawant, S Digraskar, V. Gopalkrishna

**Author notes:** Corresponding author: Sawant P.M., Enteric Viruses Group, National Institute of Virology, 20-A, Ambedkar Road, PO Box No 11, Pune 411 001, India. E mail.

## Abstract

Group A rotaviruses (RVA) are a major cause of diarrhea in neonatal calves and children. The present study examined G/P combinations and genetic characteristics of RVAs in diarrheic bovine calves in Western India. RVAs were detected in 27 samples (17.64%) with predominance of G10P[11] (51.85%), followed by previously unreported genomic constellations, G6P[14] (14.81%), and, G6P[4] (7.40%) and G10P[33] (3.70%). Sequencing and phylogenetic analysis revealed circulation of G10 (Lineage-5), G6 (Lineage-2), P[11] (Lineage-3), P[14] (proposed Lineage-8) and P[4] (Lineage-3) genotypes. The predominant G10P[11] strains were typical bovine strains and exhibited genotypic homogeneity. The rare, G10P[33] strain, had VP7 and VP4 genes of bovine origin but resemblance of VP6 gene with simian strain indicated possible reassortment between bovine and simian (SA11-like) strains. The VP6 and VP7 genes of other two rare strains, G6P[14] and G6P[4], were similar to those of bovine stains, but the VP4 was closely related to those of the human-bovine like and human strains, respectively. Additionally, in VP4 gene phylogenetic tree Indian P[14] strains constituted a closely related genetic cluster distinct from the other P[14] strains, hence Lineage-8 was proposed for them. These findings indicated that bovines could serve as source for anthropozoonotic transmission of G6P[14] strains while zooanthroponotic transmission followed by reassortment with human strain gave rise to G6P[4] strains. The observations of present study reinforce the potential of rotaviruses to cross the host-species barrier and undergo reassortant to increase genetic diversity which necessitates their continuous surveillance for development and optimization of prevention strategies against zoonotic RVAs.

## 1. Introduction

Rotaviruses are the major etiological agents responsible for acute gastroenteritis in young ones of human and animals worldwide. Rotavirus belonging to the family *Reoviridae*, is a dsRNA virus with 11 segments enclosed in triple layered capsid, and codes for six structural (VP1–VP4, VP6, and VP7) and six non structural proteins (NSP1-NSP6) (Desselberger, 2014). On the basis of sequence and antigenic variation in VP6 this virus is classified into eight groups from A to J (Crawford et al., 2017). Another two structural proteins (VP4 and VP7) bearing major antigenic determinants are used for binary classification system resulting in G (glycoprotein) and P (protease-sensitive protein) genotypes. So far, 36 G genotypes and 51 P genotypes have been characterized in humans and animals by the Rotavirus Classification Working Group (RCWG) (https://rega.kuleuven.be/cev/viralmetagenomics/virus-classification/rcwg).

Bovine rotaviruses most commonly belong to G types 6, 8 and 10 and P types [1], [5], [6] or [11] (Papp et al., 2013). In India, bovine diarrhea associated with G10 rotavirus has been seen in combination with P[11], P[6], P[14] and P[3], P[15] (Rajendran and Kang, 2014). Besides this, Indian G6 genotype has been reported in association with P[11], P[1], P[6] and P[14] genotypes (Chitambar et al. 2011). In addition to these typical G and P types, a plethora of rare genotypes with different G types (G1 to G6, G8, G10 to G12, G15, G21, G24) and P types (P[1], P[3], P[5], P[6], P[7], P[10], P[11], P[14], P[15], P[17], P[21], P[29], P[33]) have been occasionally detected in bovine diarrhea from worldwide (Kumar et al., 2018).

The combination of interspecies transmission and reassortment between RVAs of different species lead to the emergence and spread of zoonotic rotavirus strains (Desselberger, 2014). The worldwide detection including India of bovine rotavirus genomic constellations, G10P[11] and G6P[14], in neonates have raised concerns about zoonotic transmission (Iturriza-Gomara et al. 2004; Matthijnssens et al., 2009; Desselberger, 2014, Mandal et al., 2016). Besides this zooanthraponotic transmission reported in bovine (G2P[4], G1P[11] and G9P[X]) and ovine (G2P[4] and G1P[8]) species suggest complex epidemiology of rotavirus in India (Rajendran and Kang, 2014; Choudhary et al., 2017; Kumar et al., 2018). This is of great concern for developing countries like India where human-animals live in close contact and rotavirus vaccine is recently introduced in children (Ghosh et al., 2008; Malik et al., 2012). Under this scenario collecting of data on genetic diversity of bovine rotaviruses circulating in local cattle and Buffalo population will help in optimization of currently available human vaccines and formulation of future bovine vaccines. Here, the present study reports on molecular characterization of previously unreported zoonotically important G6P[14] genotypes, unusual G10P[33] and G6P[4] reassortant RVA strains from claves.

## 2. Material and methods

### 2.1. Samples

A total of 153 faecal samples were collected from diarrheic calves aged below 6 months during 2017-2019 from the Bhagyalaxshmi Dairy Farm, Manchar (80) and Govt Bull Mother Farm, Tathawade (17) in Pune district and Veterinary College, Shirwal (56) in Satara district of the Maharashtra state, Western India. A 30% faecal suspension (w/v) in phosphate buffered saline was clarified by centrifugation at 2000 X g for 10 min and supernatant was stored at −20°C till further process.

### 2.2. RVA detection, genotyping and sequencing

The total RNA was extracted from faecal suspensions by Trizol method (Invitrogen, Carlsbad, CA). Firstly, samples were screened for bovine RVA VP6 gene as per Iturriza-Gómara et al., 2002, using One Step RT-PCR kit (Qiagen, Germany) and its larger portion was amplified for phylogenetic analysis by self designed primers: VP6F 5’-CAA ATG ATA RTT ACT ATG AAY GG-3’ and VP6R 5’-ARC ATG CTT CTA ATG GAR G-3’ (product size 1079 bp). Thereafter, under identical conditions, positive samples were subjected to first round amplification of VP7 and VP4 genes as described by Isegawa et al., 1993. The samples failed in first approach were amplified using primers reported by Iturriza-Gómara et al., 2004. For amplification, 5μL denatured (97°C, 5 min) and snap chilled RNA was added in 20 µl RT-PCR mix containing 5 µl of 5x RT-PCR Buffer, 1 µl of dNTP Mix, 1 µl each Forward and Reverse primer, 1 µl of Enzyme Mix and 11 µl of Nuclease free water. The mixture was incubated at 50°C for 30 min, preheated at 95°C for 15 min, subjected to 36 cycles of 94°C for 1 min, 50°C for 1 min and 72°C for 1 min, and final incubation at 72°C for 10 min. The products were visualized in 2% agarose gel containing ethidium bromide (0.5 μg/ml) at 100 volts for about 60 minutes in 1X TAE buffer. Subsequently, amplified products were purified using agarose QIAquick Gel Purification Kit (Qiagen, Germany).

Rotavirus G and P genotyping was performed as described by Gouvea et al. 1994 and Falcone et al. 1999. Briefly, 1 µl 1:20 times diluted VP7 or VP4 template was added to 2.5 µl of 10 X PCR Buffer, 0.75 µl of 50 mM MgCl_2_, 0.5 µl of dNTPs, 0.25 µl of Taq Polymerase, 0.5 µl of G or P type primers and nuclease free water to make 25 µl reaction volume. The PCR reaction was carried out at 94°C for 4 min, then 30 cycles of 94°C for 1 min, 50°C/43°C for 2 min (G/P typing), 72°C for 2 min, and final extension at 72°C for 7 min. The amplified products were visualized as mentioned above.

All purified products of VP6 gene, and first round product of VP7 and VP4 genes were subjected to cycle sequencing using the Big Dye Terminator cycle sequencing kit v 3.1 (Applied Biosystems, Foster city, CA, USA), purified by using DyeEx 2.0 Spin Kit, (Qiagen, Germany) and sequenced on an automated Genetic analyzer ABI-PRISM3720 (Applied Biosystems). Chromatograms were analyzed in Sequencing Analysis 5.2.0 (Applied Biosystems, CA, USA). Genotypes were determined by Basic Local Alignment Search Tool (BLAST) and confirmed with RotaC 2.0 genotyping tool (Maes et al., 2009). Multiple sequence alignment of edited sequences, reference strains and cognate gene sequences available in GenBank was performed using the ClustalW algorithm. The phylogenetic trees were constructed by maximum likelihood method in MEGA v6.06 programme by using the best fit nucleotide substitution model obtained by Akaike information criterion, corrected (AICc). The best models found were GTR + G+I (VP6, VP7-G10, VP7-G6, VP4-P[33]), TN93+G+I (VP4-P[14]), HKY+I (VP4-P[11]), T93+I (VP4-P[4]). The reliability of tree was assessed by bootstrap test with 1000 replicates (Tamura et al., 2013). Lineages to the genotypes were assigned according to the literature including G10 (Komoto et al., 2016), G6 (Badaracco et al., 2013), P[11] (Badaracco et al., 2013), P[14] (Tam et al., 2014) and P[4] genotype (Tacharoenmuang et al., 2016). Nucleotide and amino acid homology was also computed using MEGA software by pairwise distance using p-distance model.

## 3. Results

### 3.1. *RVA detection, genotyping and* sequencing

In VP6 gene based screening a total of 27 (17.65%) samples were found positive for bovine RVA. The multiplex PCR was able to type G10 in 51.85%, G6 in 18.52% and P[11] in 11.11% samples. However, a large number of samples remained untypeable in G (29.63%) and P (88.89%) genotyping. Because of this, all RVA positive samples were sequenced to obtain sequence in 13 strains for VP6 gene, 21 strains for VP7 gene and 23 strains for VP4 gene. The VP7 and VP4 gene sequences were subjected for RotaC 2.0 genotyping which yielded G10 (51.85%), G6 (25.93%), P[11] (51.85%), P[14] (14.81%), P[4] (14.81%) and P[33] (3.70%) genotypes. The detected G and P type combinations included G10P[11] (51.85%), G6P[14] (14.81), G6P[4] (7.40%) and G10P[33] (3.70%). Four of the G10 and one of the G6 strains were not included in phylogenetic tree due to their partial sequences. However, their genotype were detected and included. After employing RotaC 2.0, 22.22%/14.81% samples remained non typeable in G/P typing as they failed in sequencing. The VP6, VP7 and VP4 genes sequences identified in this study were submitted to the GenBank database under the accession numbers: MT007803 to MT007815 for VP6 genes, MT007816-MT007831 for VP7 genes, MT007832-MT007851 and MT012826 for VP4 genes.

### 3.2. Sequencing and phylogenetic analysis of VP6 gene

Analysis of VP6 gene sequences showed all study strains clustering under I2 genotype in three different clusters (Fig 1a). The G6P[14] and G6P[4] strains shared identity in range of 93.3-98.8% (nucleotide) and 99.3% (amino acid) with other strains in cluster 1. Interestingly, this cluster comprised of unusual Indian bovine strain BRV133 (G3P[11]) as well as Indian porcine-bovine like strains (HP113 and HP140, G6P[13]), and human-bovine like strains [(MCS-KOL-383 (G10P[14]), N-1(G6P[14])] whose VP6 gene is believed to be derived from Indian bovine strain (Ghosh and Kobayashi, 2014; Mandal et al., 2016). The G10P[11] study strains exhibited 94.9-95.6% (nucleotide) and >99.3% (amino acid) similarity with cluster 2. In cluster three, Dai-10/SA11 cluster, G10P[33] study strain exhibited higher nucleotide identity of 97.2%, 94.7%, 93.5%, 92.9% and, 91.7%, 91.5% with Japanese human strain 12597(G8P[14]), Slovenian Roe deer strain D110-15 (G8P[14]), American human strain 2012741499, Simian strain SA11-H96 (G3P[5]), Indian bovine strain RUBV51 (G15P[11]) and Japanese bovine strain Dai-10 (G24P[33]), respectively. Surprisingly, their deduced amino acid sequences showed 100 % identities.

**Fig 1:**
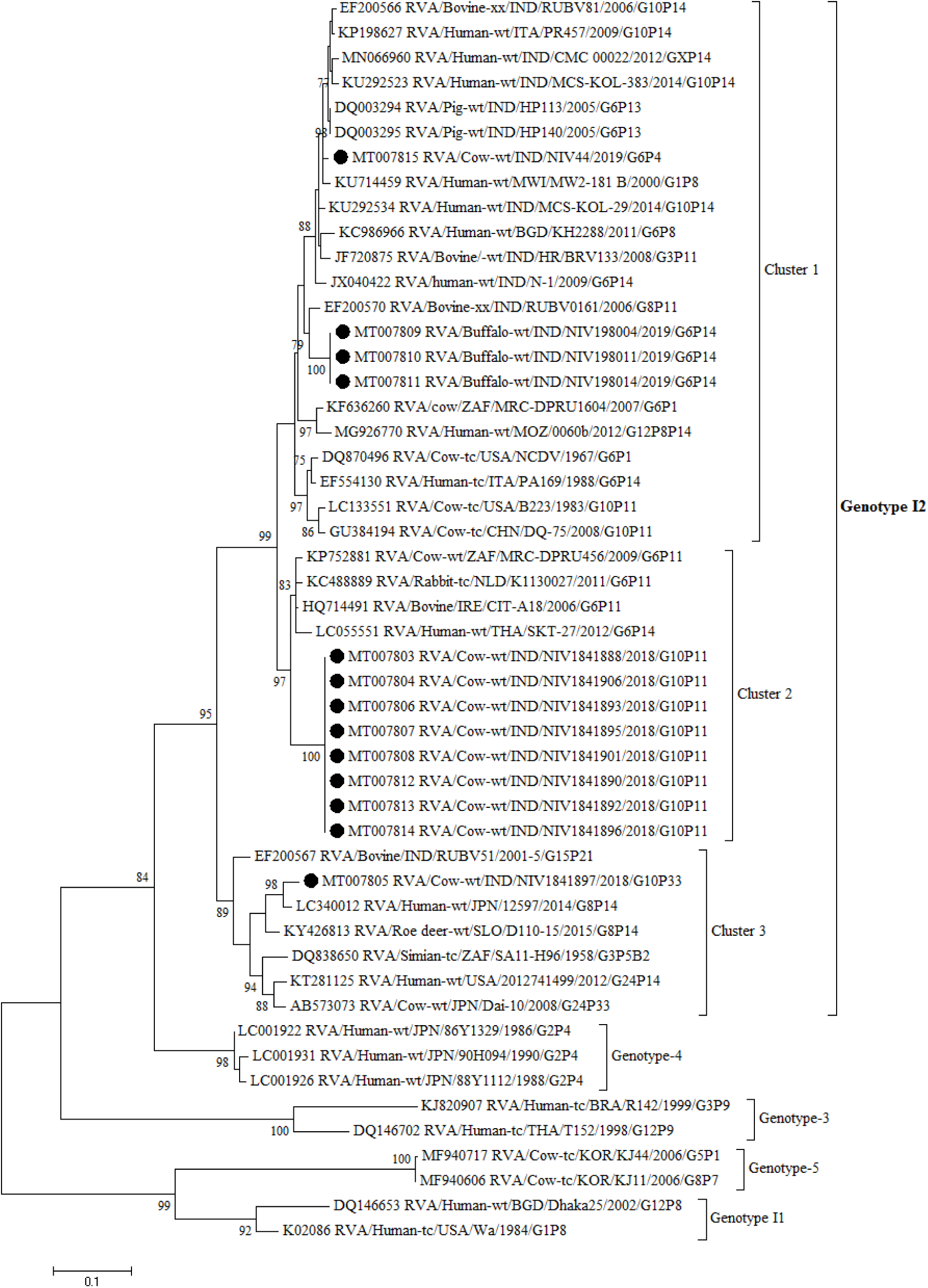
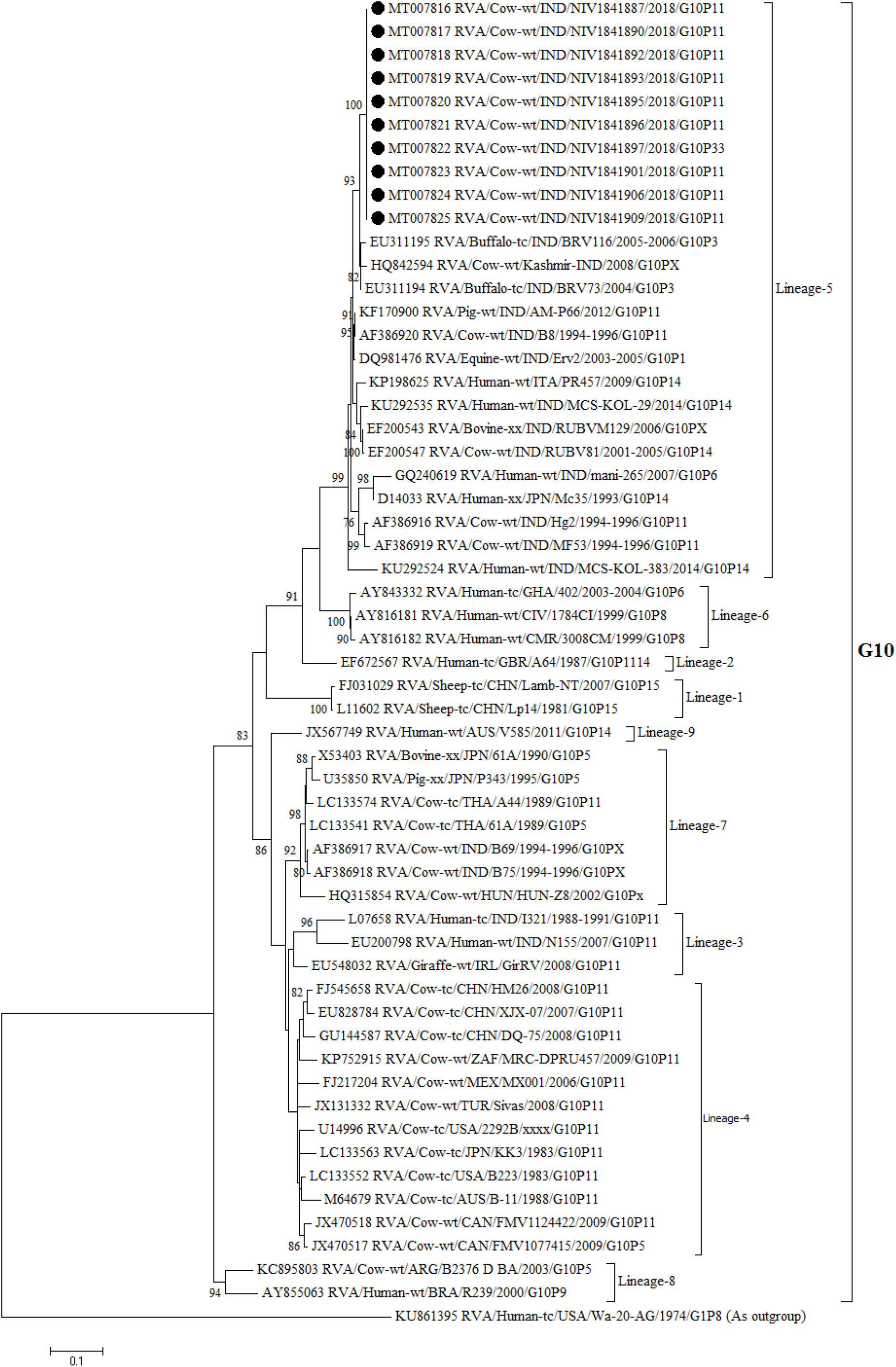
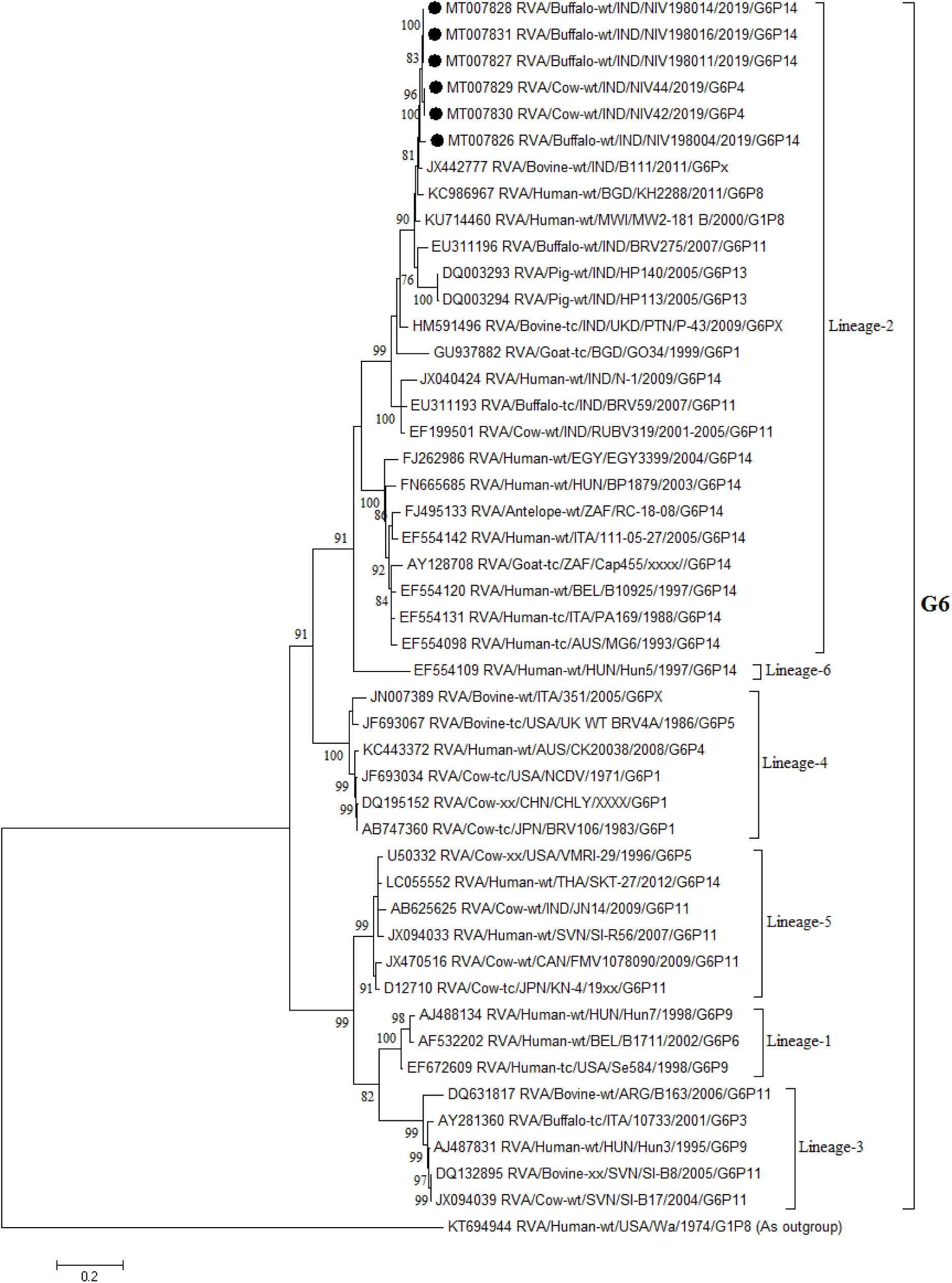
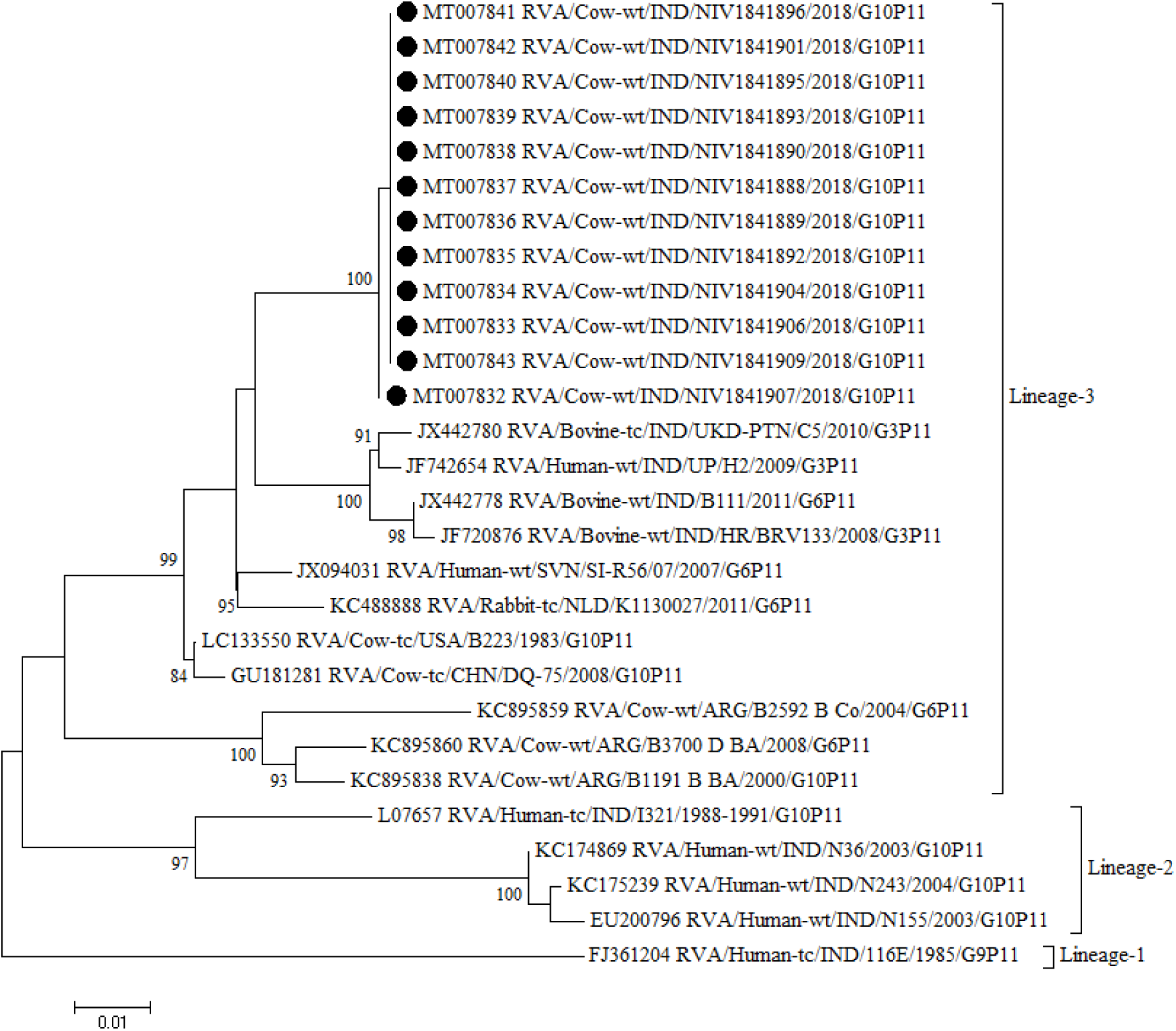
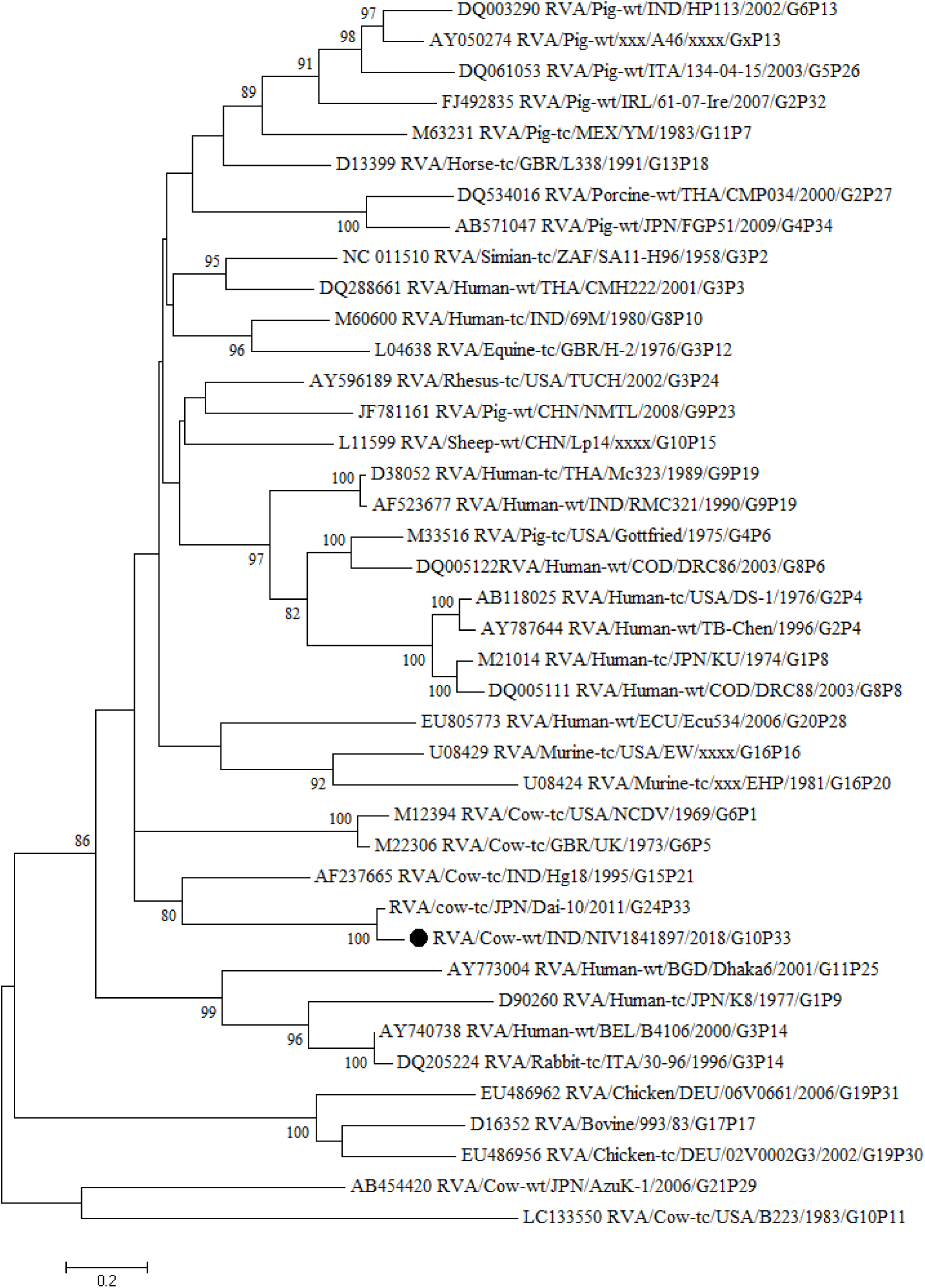
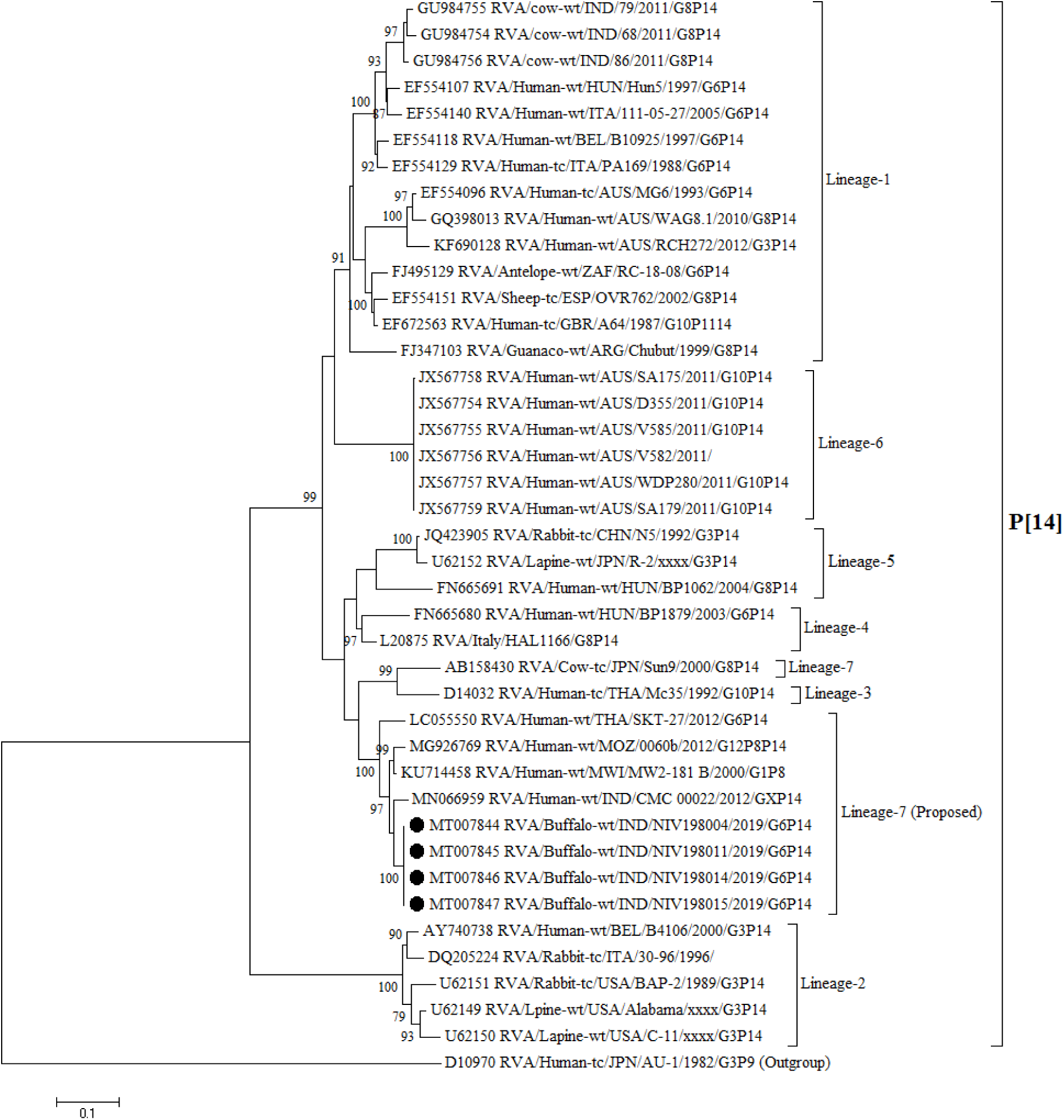
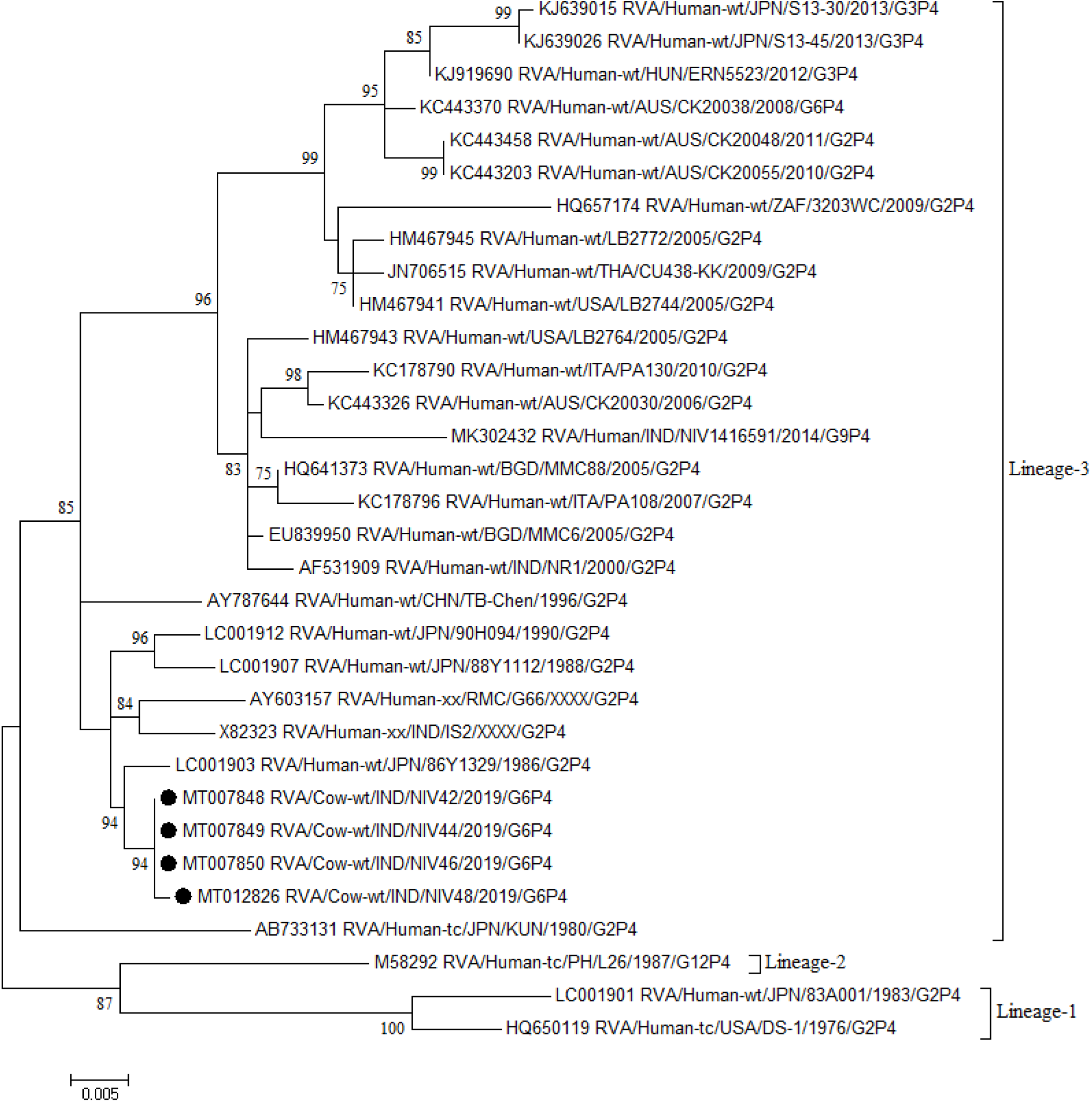
Phylogenetic trees constructed from the nucleotide sequences of the VP6 (a), G10-VP7 (b), G6-VP7 (c), P[11]-VP4 (d), P[33]-VP4 (e), P[14]-VP4 (f), P[4]-VP4 (g) genes of study strains and representative RVA strains. In all the trees, the positions of study strains are shown in black dots. Bootstrap values of <75% are not shown.

### 3.3. Sequencing and phylogenetic analysis of VP7 gene

The detected G10P[11] and G10P[33] strains grouped in Lineage-5 with 93.1-98.6% (nucleotide) and 94.7-99.5% (amino acid) identity (Fig 1b). They exhibited maximum 98.6% (nucleotide) and 99.5% (amino acid) sequence identity with Indian bovine strain BRV73 (G10P[3]). In phylogeny, earlier reported Indian G10 strains segregated in three different lineages. The G10 strains recovered in present study grouped in Lineage-5 in which majority of the Indian bovine strains (BRV73, BRV116, Hg2, B8, MF53) fall along with porcine (AM-P66), equine (Erv2) and human-bovine like strains (mani-265, MCS-KOL-29, MCS-KOL-383). The other two lineages with Indian G10 strains were Lineage-7 with bovine strains (B69 and 75) and Lineage-3 with human bovine like strains (I321 and N155). The detected G6 strains with P[14] and P[4] genotypes fall in Lineage-2 (Fig 1c). Their sequences were more than 97% (nucleotide) and 98% (amino acid) identical with each other, and 86.1-97.6% (nucleotide) and 95.3-98.8% (amino acid) identical with other Lineage-2 strains. The G6 study strains exhibited maximum identity of 96.7-97.6% (nucleotide) and >98.4% (amino acid) with that of Indian bovine strain B111 (G6P[11]) and Bangladeshi human-bovine like strain KH2288 (G6P[8]). They were grouped into G6 Lineage-2 where majority of human-bovine like G6P[14] strains reported worldwide fall along with Indian bovine strains (B111, BRV275, P-43, BRV59, RUBV319), human-bovine like strain (N-1), and porcine-bovine like strains (HP113, HP140).

### 3.4. Sequencing and phylogenetic analysis of VP4 gene

In phylogenetic analysis of reference and selected GenBank strains with P[11], P[14] and P[4] genotype segregated into three, seven and three lineages respectively. On sequence analysis, all study strains shared more than 99% identity among respective genotypes. The study strains with P[11] genotype grouped in Lineage-3, showed 96.3-97.5% (nucleotide) and >98.7% (amino acid) identity with prototype bovine strain B223 (G10P[11]), other bovine strains (DQ-75, C5, BRV133), lapine strain K1130027, and human strains (SI-R56, H2). However, RVA strains with P[11] type detected from worldwide showed no association between lineages and G type, year or location (Fig 1d). The P[33] study strain shared highest nucleotide and amino acid identity of 93.6% and 97.5% with only P[33] strain, Dai-10 detected in bovine species from Japan (Fig 1e). Presently, P[14] genotypes are known to segregate into six lineages (Tam et al., 2014). However, study strains formed distinct cluster with human-bovine like strains (MW2-181B, CMC-00022, 0060b and SKT-27) which was different from known lineages (Fig 1f). To clarify this within and between average percent nucleotide identity (PNI) for full VP4 gene sequences of known lineages and above group of strains, which shared highest nucleotide and amino acid identity (93.2%-96.9% and 97.2-99.2%) with study strains, was calculated. The within and between average PNI of known lineages ranged from 91.6-100% and 80.7-89.1% whereas for proposed lineage it was 95% and 81.3-89.5%. Therefore, this finding indicates that P[14] study strains belong to a new lineage (proposed Lineage-8) along with above mentioned human-bovine like strains. The P[4] study strains showed maximum identities in the range of 98.7-99.4% (nucleotide) and >98.50% (amino acid) with human strains from Japan (86Y1329, 90H094 and 88Y1112) and India (IS-2). All the detected P[4] strains clustered in Lineage-3 along with above mentioned strains (Fig 1g).

## 4. Discussion

Bovine group A rotaviruses play an significant role in causing gastroenteritis in young calves and children in developing countries. After extensive studies conducted on human rotaviruses under Indian National Rotavirus Surveillance Network, Indian rotavirus vaccine i.e. Rotavac, is introduced in the country. However, identification of rotaviruses in bovines which live in close proximity with human populations remains largely unexplored. Hence, the present study was conducted to explore the bovine RVA in Pune and Satara districts (Maharashtra). In the present study, out of 153 samples analyzed from diarrheic calves, 27 (17.65%) were found positive for RVA. The prevalence of rotavirus observed in this study is consistent with the studies conducted in Kashmir (15.5%) (Beg et al., 2010), Western India (14.3%) (Chitambar et al., 2011), France (15%) (Midgley et al., 2012), Tunisia (15.4%) (Hassine Zaafrane et al., 2014) and Colombia (19.7%) (Pardo-Mora et al., 2018).

Evidences from epidemiological studies worldwide indicates, G6 and G10 along with P[5] and P[11] are the most common genotypes identified in bovines. The predominance of G10P[11] combination observed in this study is in concurrence with the previous reports from India (Beg et al., 2010; Minakshi et al., 2015, Tatte et al., 2019). However, these findings are in contrast with Indian studies with G3P[11] predominance (Malik et al., 2012; Sravani et al., 2014) and reports from Iran (Pourasgari *et al*., 2016) and Ireland with G6P[5] as major genotype (Collins *et al*., 2014). Interestingly, the VP6 gene of G10P[11] study strains grouped in different cluster than prototype Bovine strains, in I2 genotype. Although, the VP4 gene of all study strains (G10P[11]) was similar to reference strain B223 (G10P[11]) their VP7 gene clustered in Lineage-5 with indigenous bovine strain BRV73 (G10P[3]), away from Lineage-4 where the prototype strain clustered. Nevertheless VP7 and VP4 genes showed similarity with Indian bovine strains but no Indian strain grouped with both genes, indicating that it might be unique strain. Such intragenotype heterogeneity among G10P[11] strains should be considered while development of bovine rotavirus vaccine, because these strains may escape the immune pressure induced by vaccines.

P[33], a rare genotype, has been described once from asymptomatic cows in Japan for rotavirus strain, Dai-10 (G24P[33]) (Abe et al., 2011). Thereafter, there were no reports on detection of P[33] genotype and to the best of our knowledge, the G10P[33] combination detected in diarrheic claves from the present study was never reported from cattle or other species. Although, the VP7 gene of G10P[33] strain was of bovine origin, its VP6 gene showed higher level of identity with Simian strain SA11-H96 and rare strains with unusual genotypes, which are distantly related to most of the bovine strains. These observations indicate that VP6 gene might have derived from reassortment events between bovine and simian (SA11-like) RVAs. The detection of these strains from different regions in Asia and Europe pointed towards wider circulation albeit at low rate of SA11-H96-like VP6 gene in ruminants.

The second predominant genotype combination found in the present study was G6P[14] (12.5%). Previously, P[14] genotype have been detected with three common bovine G types (G6, G8, G10) in artiodactyls and in humans as zoonotic strains. Out of these only G8P[14] and G10P[14] combinations are reported in cattle (Ghosh and Kobayashi, 2014). The present study, first time reports RV strains with G6P[14] combination in buffalo species. These strains clustered with human-bovine like strains under I2 genotype and G6 Lineage-2 strains with P[14] specificity, away from the G6 Lineage-4 comprising typical P[1], P[5] and P[11] bovine RVA strains from all over the World (Badaracco et al., 2013). In VP4 phylogeny, previously reported Indian bovine G8P[14] strains (BRV68, BRV79, BRV86) from Pune region and G10P[14] strains (RUBV 81) occupied Lineage-1 with Indian human-bovine like strain N-1 (G6P[14]) (Chitambar et al., 2011; Mullick et al., 2013). In contrast the study strains grouped in proposed Lineage-8 along with Indian human-bovine like GXP[14] strain (CMC-00022), and G10P[14] strains (MCS-KOL-383 and KOL-29), earlier considered in Lineage-4 (only VP8* region sequence available) (Mandal et al., 2016). The higher sequence identities of VP6, VP7 and VP4 genes and grouping in same lineages with strains previously known for interspecies transmission hypothesize zoonotic potential of bovine G6P[14] strains detected in the present study. Further, detection of G8P[14], G10P[14] and G6P[14] combinations in bovines indicates their persistence in animals, while possibility of transmission to human is supported by detection of G10P[14] and G6P[14] combinations in India. Such zoonotic transmission of P[14] genotype is possible because the VP8* region of VP4 protein interaction with type A histo blood group antigens of humans as well as bovine and porcine mucins (Tam et al., 2014).

RV P[4] genotype with G2 specificity is one of the most common types observed in humans. The combining of P[4] with different G-type VP7 genes has resulted into different G/P combinations including, G2P[4], G1P[4], G3P[4], G4P[4], G9P[4], G11P[4] and G12P[4] (L26) (Chen et al., 2008). Additionally, the sequence of only one human G6[4] strain from Australia is available, however, the G6P[4] study strain detected from bovine is first report. The close phylogenetic relationship of VP6 and VP7 genes of G6P[4] study strain with bovine strains while VP4 gene identity with typical human P[4] strains suggests that this is bovine-human reassortant strain. This strain might have evolved after infection of bovine with human rotavirus carrying P[4] genotype followed by reassortment with bovine G6 strain. These findings bolster the zooantraponotic transmission reported earlier in bovine (G2P[4], G1P[11] and G9P[X]) and ovine (G2P[4] and G1P[8]) species from India (Rajendran and Kang, 2014; Choudhary et al., 2017).

In conclusion, the present study revealed circulation of G10P[11], G6P[14], G10P[33] and G6P[4] bovine RVAs in diarrheic calves of Western India. The evidence of unusual genomic constellations in bovines, G10P[33], G6P[14] and G6P[4], exhibits the high genetic diversity of rotavirus in the reported region. The new Lineage-8 proposed in this study highlights expanding sequence diversity within the P[14] genotype, thus contributing to a better understanding of zoonotically important rotavirus genotype. Finally, presence of reassortant and zoonotic rotaviruses emphasizes their simultaneous monitoring in animals and human for the development and optimization of RV vaccines.

## Author contributions

Conception and Design, S.P.,; Performing the Experiments, Designing the Work, Acquisition, Analysis, and Interpretation of data, S.P., D.S.,; Writing the Original Manuscript, S.P.; Manuscript Revision and over all Execution: S.P.; V.G.

## Acknowledgement

The authors thank Director, ICMR-National Institute of Virology, Pune for intramural funding and constant support. We also thank Dr. Amol Hande, Farm Manager, Bhagyalaxshmi Dairy Farm, Manchar, Pune for providing clinical samples.

## References

1. Abe, M., Ito, N., Masatani, T., Nakagawa, K., Yamaoka, S., Kanamaru, Y., Suzuki, H., Shibano, K., Arashi, Y., Sugiyama, M., 2011. Whole genome characterization of new bovine rotavirus G21P[29] and G24P[33] strains provides evidence for interspecies transmission. J. Gen. Virol. 92(Pt 4), 952–960. https://doi.org/10.1099/vir.0.028175-0.

2. Badaracco, A., Garaicoechea, L., Matthijnssens, J., Louge Uriarte, E., Odeón, A., Bilbao, G., Fernandez, F., Parra, G.I., Parreño, V., 2013. Phylogenetic analyses of typical bovine rotavirus genotypes G6, G10, P[5] and P[11] circulating in Argentinean beef and dairy herds. Infect. Genet. Evol. 18, 18–30. https://doi.org/10.1016/j.meegid.2013.04.023.

3. Beg, S.A., Wani, I., Hussain, Bhat, M.A., 2010. Determination of G and P type diversity of group A rotaviruses in faecal samples of diarrhoeic calves in Kashmir. Indian J. Appl. Microbiol. 51(5), 595–599. https://doi.org/10.1111/j.1472-765X.2010.02944.x.

4. Chen, Y., Wen, Y., Liu, X., Xiong, X., Cao, Z., Zhao, Q., Yu, Y., Yin, X., Li, C., Fan, Y., 2008. Full genomic analysis of human rotavirus strain TB-Chen isolated in China. Virol. 375(2), 361–373. https://doi.org/10.1016/j.virol.2008.01.003.

5. Chitambar, S.D., Arora, R., Kolpe, A.B., Yadav, M.M., Raut, C.G., 2011. Molecular characterization of unusual bovinegroup A rotavirus G8P[14] strains identified in western India: Emergence of P[14] genotype. Vet. Microbiol. 148, 384–388. https://doi.org/10.1016/j.vetmic.2010.08.027.

6. Choudhary, P., Minakshi, P., Ranjan, K., Basanti, B., 2017. Zooanthroponotic transmission of rotavirus in Haryana State of Northern India. Acta. Virol. 61(1), 77–85. https://doi.org/10.4149/av_2017_01_77.

7. Collins, P.J., Mulherin, E., Cashman, O., Lennon, G., Gunn, L., O’Shea, H., Fanning, S., 2014. Detection and characterization of bovine rotavirus in Ireland from 2006-2008. Irish Vet. J. 67(1), 13. https://doi.org/10.1186/2046-0481-67-13.

8. Crawford, S.E., Ramani, S., Tate, J.E., Parashar, U.D., Svensson, L., Hagbom, M., Franco, M.A., Greenberg, H.B., O’Ryan, M., Kang, G., Desselberger, U., Estes, M.K., 2017. Rotavirus infection. Nat. Rev. Dis. Primers 3, 17083. https://doi.org/10.1038/nrdp.2017.83.

9. Desselberger, U., 2014. Rotaviruses. Virus Res. 190, 75–96. https://doi.org/10.1016/j.virusres.2014.06.016.

10. Falcone, E., Tarantino, M., Trani, L.D., Cordioli, P., Lavazza, A., Tollis, M., 1999. Determination of G and P sero-types in Italy using PCR. J. Clin. Microbiol. 37, 3879–3882.

11. Ghosh, S., Kobayashi, N., 2014. Exotic rotaviruses in animals and rotaviruses in exotic animals. Virus dis. 25(2), 158–172. https://doi.org/10.1007/s13337-014-0194-z.

12. Gouvea, V., Satos, N., Timentsky, M.R., 1994. Identification of bovine and porcine rotavirus G types by PCR. J. Clin. Microbiol. 32, 1338–1340.

13. Hassine-Zaafrane, I., Salem, B., Sdiri-Loulizi, K., Kaplan, J., Bouslama, L., Aouni, Z., Sakly, N., Pothier, P., Aouni, M., Ambert-Balay, K., 2014. Distribution of G (VP7) and P (VP4) genotypes of group A bovine rotaviruses from Tunisian calves with diarrhoea. J. Appl. Microbiol. 116, 1387—1395. https://doi.org/10.1111/jam.12469.

14. Isegawa, Y., Nakagomi, O., Nakagomi, T., Ishida, S., Uesugi, S., Ueda, S., 1993. Determination of bovine rotavirus G and P serotypes by polymerase chain reaction. Mol. Cell Probes 7(4), 277–284.

15. Iturriza Gómara, M., Wong, C., Blome, S., Desselberger, U., Gray, J., 2002. Molecular characterization of VP6 genes of human rotavirus isolates: correlation of genogroups with subgroups and evidence of independent segregation. J. Virol. 76(13), 6596–601. https://doi.org/10.1128/jvi.76.13.6596-6601.2002.

16. Iturriza-Gómara, M., Kang, G., Gray, J., 2004. Rotavirus genotyping: keeping up with an evolving population of human rotaviruses. J. Clin. Virol. 31(4), 259–265. https://doi.org/10.1016/j.jcv.2004.04.009.

17. Komoto, S., Pongsuwanna, Y., Tacharoenmuang, R., Guntapong, R., Ide, T., Higo-Moriguchi, K. Tsuji, T., Yoshikawa, T., Taniguchi, K., 2016. Whole genomic analysis of bovine group A rotavirus strains A5-15 and A5-13 provides evidence for close evolutionary relationship with human rotaviruses. Vet. Microbiol. 195, 37–57. https://doi.org/10.1016/j.vetmic.2016.09.003.

18. Kumar, N., Malik, Y.P.S., Sharma, K., Dhama, K., Ghosh, S., Bányai, K., Kobayashi, N., Singh, R.K., 2018. Molecular characterization of unusual bovine rotavirus A strains having high genetic relatedness with human rotavirus: evidence for zoo-anthroponotic transmission. Zoonoses Public Health 65, 431–442. https://doi.org/10.1111/zph.12452.

19. Maes, P., Matthijnssens, J., Rahman, M., VanRanst, M., 2009. RotaC: a web-based tool for the complete genome classification of group A rotaviruses. BMC Microbiol. 9, 238. https://doi.org/10.1186/1471-2180-9-238.

20. Malik, Y.P.S., Sharma, K., Vaid, N., Chakravarti, S., Chandrashekar, K.M., Basera, S.S., Singh, R., Minakshi, Prasad G., Gulati, B.R., Bhilegaonkar, K.N., Pandey, A.B., 2012. Frequency of group A rotavirus with mixed G and P genotypes in bovines: predominance of G3 genotype and its emergence in combination with G8/G10 types. J. Vet. Sci. 13(3), 271–278. https://doi.org/10.4142/jvs.2012.13.3.271.

21. Mandal, P., Mullick, S., Nayak, M.K., Mukherjee. A., Ganguly. N., Niyogi. P., Panda. S., Chawla-Sarkar. M., 2016. Complete genotyping of unusual species A rotavirus G12P[11] and G10P[14] isolates and evidence of frequent in vivo reassortment among the rotaviruses detected in children with diarrhea in Kolkata, India, during 2014. Arch. Virol. 161(10), 2773–2785. https://doi.org/10.1007/s00705-016-2969-6.

22. Matthijnssens, J., Potgieter, C.A., Ciarlet, M., Parreño, V., Martella, V., Bányai, K., Garaicoechea, L., Palombo, E.A., Novo, L., Zeller, M., Arista, S., Gerna, G., Rahman, M., Van Ranst, M., 2009. Are human P[14] rotavirus strains the result of interspecies transmissions from sheep or other ungulates that belong to the mammalian order Artiodactyla? J. Virol. 83, 2917–2929. https://doi.org/10.1007/s00705-011-1006-z.

23. Midgley, S.E., Bányai, K., Buesa, J., Halaihel, N., Hjulsager, C.K., Jakab, F., Kaplon, J., Larsen, L.E., Monini, M., Poljšak-Prijatelj, M., Pothier, P., Ruggeri, F.M., Steyer, A., Koopmans, M., Böttiger, B., 2012. Diversity and zoonotic potential of rotaviruses in swine and cattle across Europe. Vet. Microbiol. 156(3-4), 238–245. https://doi.org/10.1016/j.vetmic.2011.10.027.

24. Minakshi, P., Ranjan, K., Pandey, N., Dahiya, S., Khurana, S., Bhardwaj, N., Bhardwaj, S., Prasad, G., 2015. Predominance of G10 genotype of rotavirus in diarrheic buffalo calves: a potential threat for animal to human zoonotic transmission. Adv. Anim. Vet. Sci. 3(1s), 16–21. http://dx.doi.org/10.14737/journal.aavs/2015/3.1s.16.21.

25. Mullick, S., Mukherjee, A., Ghosh, S., Pazhani, G.P., Sur, D., Manna, B., Nataro, J.P., Levine, M.M., Ramamurthy, T., Chawla-Sarkar, M., 2013. Genomic analysis of human rotavirus strains G6P[14] and G11P[25] isolated from Kolkata in 2009 reveals interspecies transmission and complex reassortment events. Infect. Genet. Evol. 14, 15–21. https://doi.org/10.1371/journal.pone.0112970.

26. Papp, H., László, B., Jakab, F., Ganesh, B., De Grazia, S., Matthijnssens, J., Ciarlet, M., Martella, V., Bányai, K., 2013. Review of group A rotavirus strains reported in swine and cattle. Vet. Microbiol. 165(3-4), 190–199. https://doi.org/10.1016/j.vetmic.2013.03.020.

27. Pardo-Mora, D., 2018. Molecular characterization of rotaviruses isolated from calves with bovine neonatal diarrhea in Colombia. Infectio. 22(2), 99–104. https://doi.org/10.22354/in.v22i2.715.

28. Park, S., Matthijnssens, J., Saif, L.J., Kim, H.J., Park, J.G., Alfajaro, M.M., Kim, D.S., Son, K.Y., Yang, D.K., Hyun, B.H., Kang, M.I., Cho, K.O., 2011. Reassortment among bovine, porcine and human rotavirus strains results in G8P[7] and G6P[7] strains isolated from cattle in South Korea. Vet. Microbiol. 152(1-2), 55–66. https://doi.org/10.1186/1297-9716-44-88.

29. Pourasgari, F., Kaplon, J., Karimi-Naghlani, S., Fremy, C., Otarod, V., Ambert-Balay, K., Mirjalili, A., Pothier, P., 2016. The molecular epidemiology of bovine rotaviruses circulating in Iran: a two-year study. Arch. Virol. 161(12), 3483–3494. https://doi.org/10.1007/s00705-016-3051-0.

30. Rajendran, P., Kang, G., 2014. Molecular epidemiology of rotavirus in children and animals and characterization of an unusual G10P[15] strain associated with bovine diarrhea in south India. Vaccine 32 Suppl 1, A89–94. https://doi.org/10.1016/j.vaccine.2014.03.026.

31. Sravani, G.V.D., Kaur, G., Chandra, M., Kaur, S.G., Chandra, M., Dwivedi, P.N., Dwivedi, P. N., 2014. P and G genotyping of bovine rotavirus and its prevalence in and G genotyping of bovine rotavirus and its prevalence in neonatal calves. Veterinarski Arhiv. 84 (5), 475–484.

32. Tacharoenmuang, R., Komoto, S., Guntapong, R., Ide, T., Sinchai, P., Upachai, S., Yoshikawa, T., Tharmaphornpilas, P., Sangkitporn, S., Taniguchi, K., 2016. Full Genome Characterization of Novel DS-1-Like G8P[8] Rotavirus Strains that Have Emerged in Thailand: Reassortment of Bovine and Human Rotavirus Gene Segments in Emerging DS-1-Like Intergenogroup Reassortant Strains. PLoS One 11(11), e0165826. https://doi.org/10.1371/journal.pone.0165826.

33. Tam, K.I., Roy, S., Esona, M.D., Jones, S., Sobers, S., Morris-Glasgow, V., Rey-Benito, G., Gentsch, J.R., Bowen, M.D., 2014. Full genomic characterization of a novel genotype combination, G4P[14], of a human rotavirus strain from Barbados. Infect. Genet. Evol. 28, 524–529. https://doi.org/10.1016/j.meegid.2014.09.020.

34. Tamura, K., Stecher, G., Peterson, D., Filipski, A., Kumar, S., 2013. MEGA6: Molecular Evolutionary Genetics Analysis version 6.0. Mol. Biol. Evol. 30(12), 2725–2729. https://doi.org/10.1093/molbev/mst197.

35. Tatte, V.S., Jadhav, M., Ingle, V.C., Gopalkrishna, V., 2019. Molecular characterization of group A rotavirus (RVA) strains detected in bovine and porcine species: Circulation of unusual rotavirus strains. A study from western, India. Acta. Virol. 63(1), 103–110. https://doi.org/10.4149/av_2019_113.

